# Exosomal transmission of α-synuclein initiates Parkinson’s disease-like pathology

**DOI:** 10.1101/2021.05.10.443522

**Authors:** Jason Howitt, Ley Hian Low, Michelle Bouman, Jeremy Yang, Sarah Gordon, Andrew Hill, Seong-Seng Tan

## Abstract

The intercellular transmission of α-synuclein is proposed to be involved in the pathogenesis of Parkinson’s disease (PD)^1^. Supporting this transmission hypothesis, α-synuclein has been identified to act as a prion^2^ and has been found in cerebrospinal fluid (CSF), blood, and saliva^3^. However, the mechanism required to initiate the pathogenic spread of α-synuclein in the body that results in PD remains unclear. Here we identify a mechanism for the loading of α-synuclein into exosomes, resulting in prion-like transmission that is mediated by the ubiquitin ligase activator, Ndfip1. Risk factors associated with PD, including metal toxicity, lysosome dysfunction and pesticide exposure,^4^ stimulate the upregulation of Ndfip1 and increase the loading of α-synuclein into exosomes. We used this pathway to promote the loading of endogenous α-synuclein into exosomes, that when intranasally delivered to either wild type or M83 transgenic mice, resulted in Parkinson’s-like pathology including motor impairments and brain amyloids. Recipient cells require α-synuclein for pathogenesis, as delivery of exosomes containing α-synuclein to Snca knockout mice did not generate brain amyloids or Parkinson’s-like motor impairments. Our results demonstrate a novel mechanism to initiate the prion-like spread of α-synuclein in exosomes, following exposure to risk factors for PD.

---

Parkinson’s disease is the most common neurological movement disorder and is now identified as the fastest growing neurodegenerative disease worldwide^5^. Although genetic studies have identified important pathways in the disease pathology, approximately 90% of all PD cases are idiopathic in nature^6^, highlighting our limited understanding for how the disease process is initiated. Strong evidence implicates α-synuclein in the pathogenesis of PD, with brain amyloids containing the protein and genetic studies identifying mutations and multiplications in the SNCA gene from familial PD patients. Extracellular α-synuclein has been identified as a potential agent for the transmission of PD pathology from diseased to healthy cells in the body^7^. This transmission hypothesis is supported by multiple lines of evidence including: post mortem studies of PD pathology^8^, identification of α-synuclein aggregates in grafted fetal dopaminergic neurons in the brains of PD patients^9^, and animal studies where synthetic misfolded α-synuclein can ‘seed’ neuronal α-synuclein aggregation in the brain^10^. The mechanism for the secretion of α-synuclein from the cell remains controversial, with passive release of the protein being identified along with a directed pathway, where a small fraction of α-synuclein is associated with extracellular vesicles called exosomes^11^. Significantly, the lipid environment of exosomes can promote the misfolding of α-synuclein^12^ and exosomes derived from PD patients have been shown to transmit pathology to neurons *in vitro*^13^, indicating a potential route of disease transmission. As such, the mechanism of α-synuclein loading and subsequent trafficking within exosomes is of significant interest for understanding PD pathogenesis.

## Ndfip1 mediates the loading of α-synuclein into exosomes

To identify a mechanism for the loading of α-synuclein into exosomes we investigated the Nedd4-1 ubiquitin ligase pathway which has been shown to promote α-synuclein degradation via the endosomal-lysosomal system^14^. Nedd4-1 can be activated by its adaptor Ndfip1, a protein that localises to endosomes and acts as a scaffold for the recruitment of target proteins for ubiquitylation^15,16^. Ndfip1 has been identified to be upregulated in surviving neurons in the substantia nigra of PD patients in association with increased iron concentrations^17^, indicating a potential role for the protein in the disease process. To search for a direct interaction between α-synuclein and Ndfip1, we used immunoprecipitation and bimolecular fluorescent complementation (BiFC) assays. Immunoprecipitation of Ndfip1 from the mouse brain resulted in co-precipitation of α-synuclein, demonstrating an interaction between the two proteins (Fig. 1a). In cell culture, an interaction between Ndfip1 and either α-synuclein or a familial α-synuclein mutant, A53T, was found when co-expressed (Fig. 1b). BiFC assays identified this interaction to occur on both early (Rab5) and recycling endosomes (Rab11) in the cell (Fig. 1c, d). Given that Ndfip1 can activate Nedd4 ubiquitin ligases, we investigated if this complex could result in the ubiquitylation of α-synuclein. Indeed, denaturing ubiquitylation assays indicated that expression of Ndfip1/α-synuclein/Nedd4-1 could result in both mono and poly-ubiquitylation of α-synuclein (Fig. 1e). To determine the location of ubiquitylated-α-synuclein in the cell, BiFC assays were performed, which identified the conjugated-protein to be predominantly localised to late and recycling endosomes in the cell, represented by Rab7, 9 and 11 (Fig. 1f, g).

**Fig. 1.**
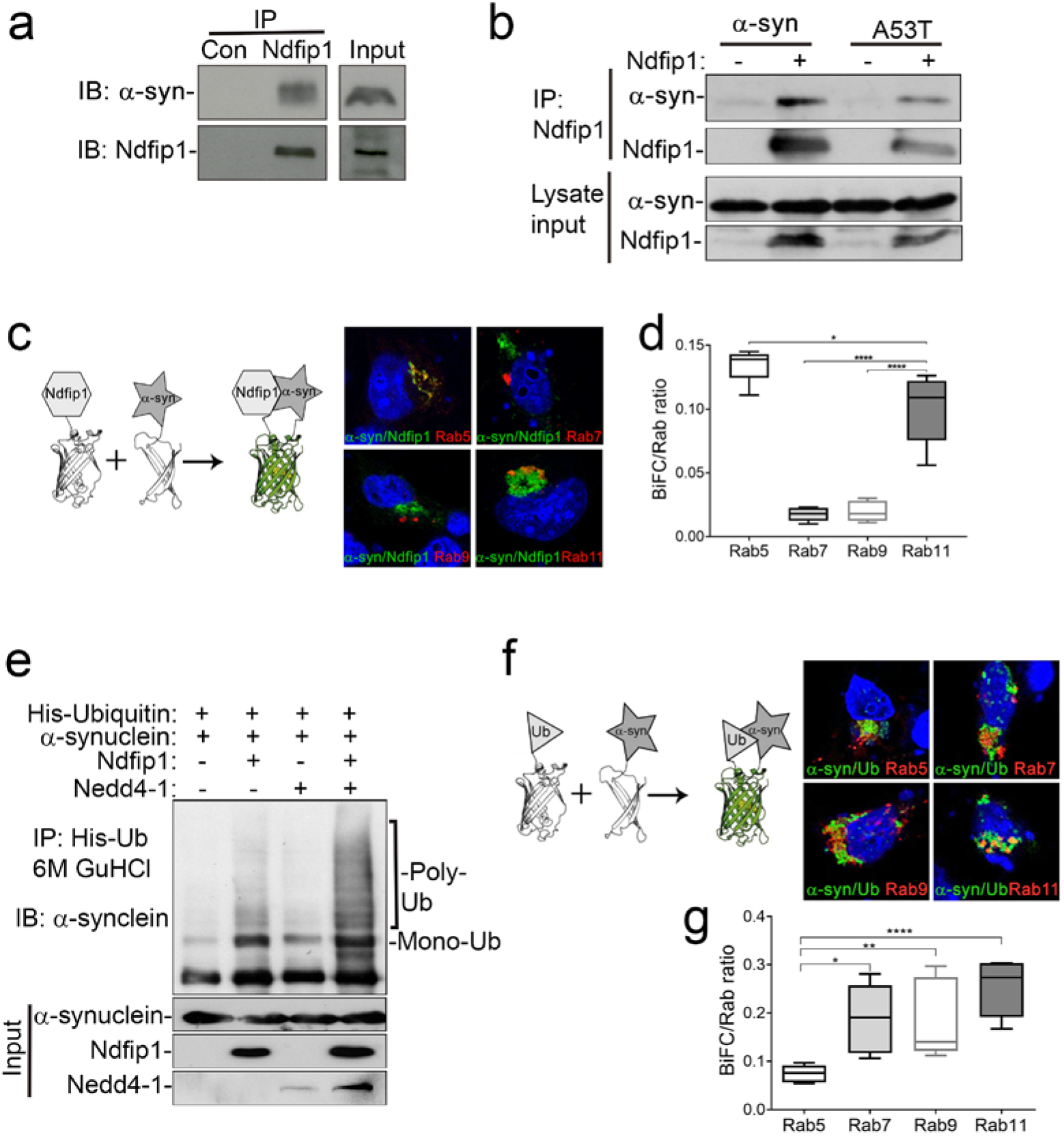
Ndfip1 interacts with and mediates the ubiquitylation of α-synuclein. **a**, Immunoprecipitation of Ndfip1 from the mouse cortex results in co-precipitation of α-synuclein. **b**, Overexpression of either α-synuclein or a familial PD mutant A53T in HEK293T cells results in co-precipitation with Ndfip1 pulldown. **c, d**, BiFC was used to visualise the location of interactions between Ndfip1 and α-synuclein. To quantify the location of the interaction between Ndfip1 and α-synuclein, Rab-GTPases were used as endosomal markers. Manders coefficients were calculated between ubiquitinated-α-synuclein and each of the Rab proteins following image deconvolution. A significantly increased localisation of Ndfip1 and α-synuclein was observed on Rab5 and Rab11 containing endosomes. **e**, Co-expression of Ndfip1, Nedd4-1 and α-synuclein in HEK293 cells results in the mono and poly-ubiquitylation of α-synuclein. HEK293 cells contain low amounts of endogenous Ndfip1 and Nedd4-1 allowing for minimal ubiquitylation of α-synuclein when only individual proteins are overexpressed. **f**, BiFC was used to visualise the location of ubiquitylated-α-synuclein in combination with Rab-GTPases. Manders coefficients were calculated between ubiquitylated-α-synuclein and each of the Rab proteins following image deconvolution. A significantly increased localisation of ubiquitylated-α-synuclein was observed on late endosomal (Rab7 and 9) and recycling endosomal pathways (Rab11). n = 8 for each Rab analysed, data represents the mean ± s.e.m. by t-test **P < 0.01, ***P < 0.001 and ****P <0.0001.

Both Rab7 and Rab11, in conjunction with the endosomal sorting complex required for transport (ESCRT), play an important role in sorting ubiquitylated proteins into exosome biogenesis pathways^18,19^. In addition, Ndfip1 is known to load specific cargo proteins into exosomes^20-22^. We therefore investigated if Ndfip1 could promote exosomal loading of α-synuclein. Exosomes were purified from HEK293T cells overexpressing α-synuclein, with or without Ndfip1 co-expression, and assayed using Western blotting. Both α-synuclein and an A53T mutant α-synuclein were loaded into exosomes in the presence of Ndfip1, in contrast, there was minimal loading of either α-synuclein or the A53T mutant into exosomes without co-expression of Ndfip1 (Fig. 2a, b). Expression of Nedd4-1 (without Ndfip1) was not able to load α-synuclein into exosomes, indicating the important role Ndfip1 plays in this mechanistic pathway (Extended data Fig. 1a). Inhibition of exosome secretion by inactivation of neutral sphingomyelinase 2 with GW4869, limited the release of extracellular α-synuclein (Extended data Fig. 1b). As α-synuclein is known to bind lipids, we investigated if the protein was carried within the lumen of exosomes or bound to the lipid envelope of the vesicle. Using proteolytic digestion we confirmed that the majority α-synuclein was contained in the lumen of exosomes and not on the surface of vesicles (Fig. 2c). Next we assayed if this pathway for loading α-synuclein into exosomes could deliver α-synuclein to recipient cells. Exosomes derived from HEK293T cells expressing both Ndfip1 and GFP-labelled α-synuclein were able to transmit α-synuclein-GFP to recipient HEK293T cells, whilst expression of α-synuclein-GFP without Ndfip1 did not result in exosomal transmission when incubated with recipient cells (Fig. 2d).

**Fig. 2.**
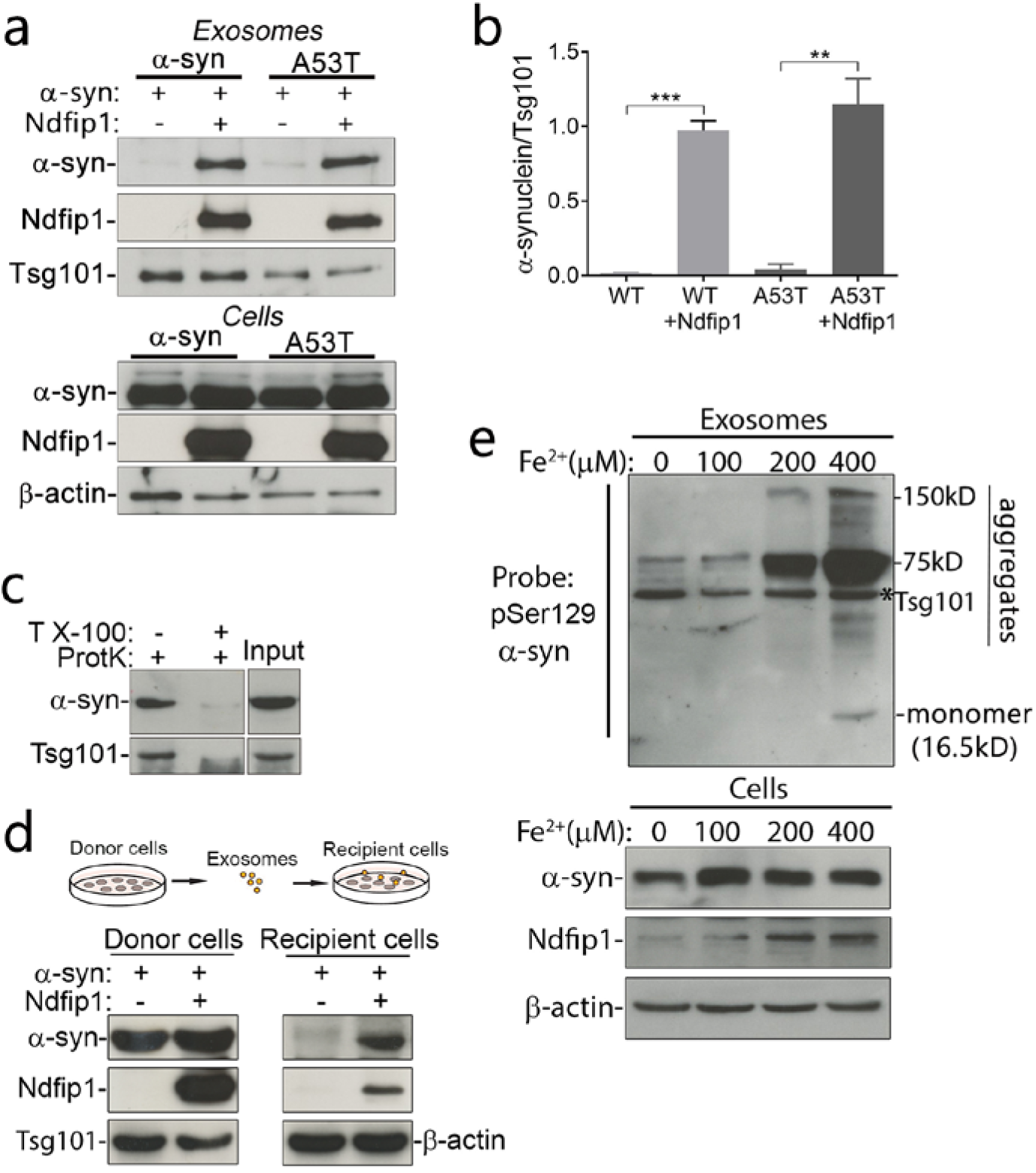
Ndfip1 is required for packaging α-synuclein into exosomes. **a, b**, Co-expression of either α-synuclein or A53T mutant α-synuclein with Ndfip1 resulted in the packaging of α-synuclein into exosomes. Quantification of Western blots showed a significant increase in exosomal α-synuclein when Ndfip1 was co-expressed in donor cells. Data are the mean ± s.e.m. by t-test * P < 0.05, ***P < 0.01, n = 3 independent experiments. **c**, Alpha-synuclein contained in the lumen of exosomes is resistant to proteinase K treatment, but sensitive to degradation after treatment with Triton X-100 (TX-100) to permeabilize exosomes. **d**, Harvested exosomes containing α-synuclein are able to transmit exogenous α-synuclein to recipient cells. **e**, Western blot showing that increasing concentrations of the PD risk factor, iron, is able to upregulate endogenous Ndfip1 in cells, resulting in the exosomal loading of endogenous α-synuclein into exosomes. Both monomer and aggregated forms of α-synuclein were able to be detected in exosomes when probed with pSer129 α-synuclein antibody. * exosomal loading control protein Tsg101. Data in **b** represents the mean ± s.e.m. by t-test **P < 0.01, ***P < 0.001.

Ndfip1 is a stress response protein that can be upregulated after various events including traumatic brain injury^23^, metal toxicity^24,25^, inflammation^26^ and DNA damage^27^. These cellular stress events overlap with multiple risk factors associated with PD^4^, we therefore investigated if risk factors associated with PD could result in the upregulation of Ndfip1 and the subsequent loading of α-synuclein into exosomes. Ndfip1 was upregulated in response to iron exposure, lysosomal inhibition and the pesticide rotenone (Fig 2e, Extended data Fig. 2a, b). To test if PD associated risk factors could promote the loading of α-synuclein into exosomes, we exposed LN18 cells containing endogenous α-synuclein and Ndfip1 to either iron toxicity or lysosome inhibition. With increasing concentrations of either iron or chloroquine (a lysosome inhibitor) we observed upregulation of Ndfip1 and the subsequent loading of endogenous α-synuclein into exosomes. Significantly, we identified both monomer and aggregated adducts of endogenous α-synuclein in exosomes when probed with a pSer129 α-synuclein antibody that detects aggregated forms of the protein (Fig. 2e and Extended data Fig. 2a). Longitudinal analysis of exosomes over time showed increased amounts of aggregated α-synuclein, suggesting that the exosome micro-environment can promote the progressive aggregation of α-synuclein as previously reported^12^ (Extended data Fig. 2c).

## Administration of exosomes containing α-synuclein results in Parkinson’s-like pathology

While *in vitro* studies have demonstrated intercellular transmission of α-synuclein via exosomes, it is unknown if this can occur *in vivo* with pathological consequences in the brain. We previously showed that exosomes delivered to the nasal passage can cross into the brain and deliver functional protein to multiple regions^22^. To test if endogenous α-synuclein from LN18 cells can be pathogenic, cells were stressed by iron exposure to upregulate Ndfip1 resulting in the loading of endogenous α-synuclein into exosomes. These exosomes were purified and delivered intranasally to M83 transgenic mice (which express the human familial PD mutant A53T, and are used to observe accelerated neurodegeneration phenotypes). M83 mice can develop motor impairments and brain amyloids containing α-synuclein at approximately 8-16 months of age^28^. To avoid this later age period, we delivered equivalent amounts of exosomes derived from both control (cells with no iron exposure) and iron stressed cells (termed ‘α-synuclein exosomes’) to mice from 2-6 months of age and monitored them for behavioural changes (Fig. 3a, Extended data Fig. 3). As PD is primarily diagnosed by motor symptoms we focussed on behavioural testing of motor function. At approximately 3.5 months of age the M83 mice given α-synuclein exosomes showed hind limb clasping which was not observed in mice given control exosomes (Fig 3b). Further behavioural testing investigating motor function were performed including a pole test, beam test and DigiGait analysis to detect motor abnormalities. At 4 months of age M83 mice given α-synuclein exosomes were observed to fail on both the pole and beam tests that require hind limb motor function to perform correctly. By 6 months of age over half of the M83 mice given α-synuclein exosomes could not perform on either the pole or beam test (Fig.3d, f, Extended data video 1). DigiGait analysis at 6 months revealed that the M83 mice given α-synuclein exosomes could not complete treadmill running at higher speeds and showed significantly increased paw angle variability, indicating less stride-to-stride consistency in the orientation of each paw placement (Fig. 3g, h). Other gait parameters, including stride length, stance width and percentage shared stance were not found to be altered, and all mice showed normal weight gain during the time period of exosome delivery (Extended data Fig. 4). M83 mice given control exosomes showed no change in motor function and were able to complete all behavioural testing at 6 months of age (Fig. 3d, f, g, h). To investigate if iron toxicity by itself could induce PD-like pathology we spiked control exosomes with iron and performed nasal delivery to M83 mice. These mice did not develop any motor function abnormalities in comparison to mice given α-synuclein exosomes (Fig. 3c, f)

Transgenic M83 mice do not represent a physiological level of α-synuclein in the body, as such we also investigated motor impairments in wild-type (WT) mice treated with α-synuclein exosomes. WT mice treated with α-synuclein exosomes were observed to show motor function impairments on the pole test (the most difficult motor test) at approximately 5 months of age, indicating a slower disease trajectory compared to the M83 mice (Extended data Fig. 3b). At 6 months of age WT mice given α-synuclein exosomes showed hind limb motor function failure on both the pole and beam test (Fig. 3c-f, Extended data video 2). DigiGait analysis showed a similar phenotype to the M83 mice given α-synuclein exosomes, with failure at high speed running and increased paw angle variability observed across treadmill speeds (Fig. 3g, h). WT mice given control exosomes showed no behavioural deficits in any motor function test. These experiments demonstrate that α-synuclein exosomes harvested from stressed cells can be pathogenic with accompanying signs of Parkinson’s-like pathology when administered to both WT and M83 mice.

**Fig. 3.**
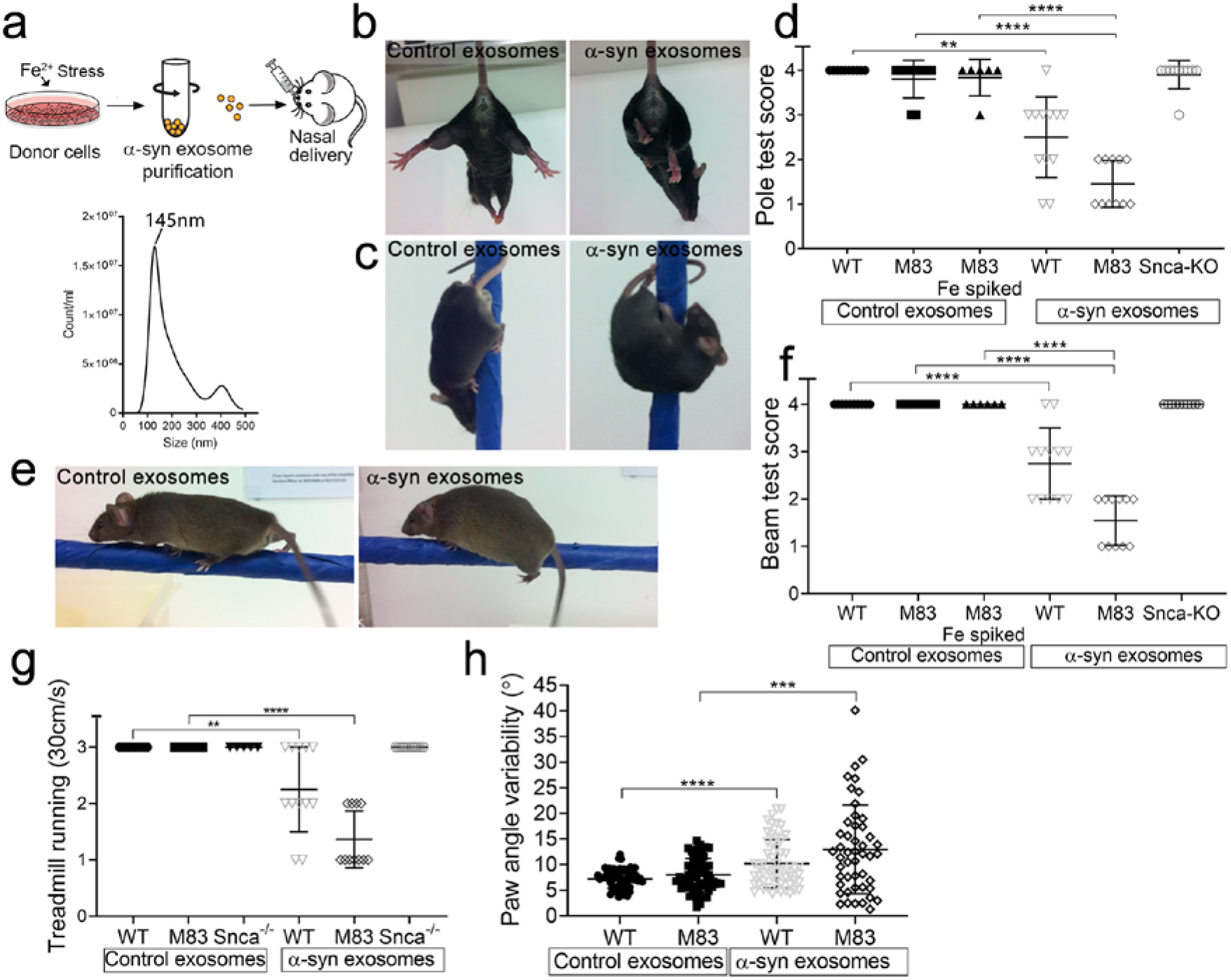
Intranasal delivery of α-synuclein containing exosomes results in motor deficits in both M83 transgenic and wild-type (WT) mice, but not Snca^>-/-^mice. **a**, Schematic for the nasal delivery of exosomes to mice and nano-particle tracking analysis (NTA) of exosome size. **b**, After delivery of α-synuclein exosomes both WT and M83 mice showed hind limb clasping (M83 mouse shown). **c**, Delivery of α-synuclein exosomes to both WT and M83 mice caused a loss in hind limb mobility, resulting in them being unable to turn on a pole test (WT mouse shown). Mice given control exosomes, or exosomes spiked with iron were able to complete the test. **d**, Quantification of pole test at six months of age showed a significantly decreased ability to turn on the pole for both WT and M83 mice after delivery of α-synuclein exosomes, delivery of α-synuclein exosomes to Snca^-/-^mice did not result in hind limb deficits. **e**, Delivery of α-synuclein exosomes to both WT and M83 mice resulted in foot faults and freezing on a beam test (WT mouse shown). Mice given control exosomes, or exosomes spiked with iron were able to complete the test with no faults. **f**, Quantification of foot faults on the beam test at six months of age, delivery of α-synuclein exosomes to Snca^-/-^mice did not result in motor deficits. **g**, Analysis of mice running on a DigiGait treadmill showed that both WT and M83 mice after delivery of α-synuclein exosomes could not complete a running test at a speed of 30cm/s. **h**, Digigait analysis of mouse treadmill running showed a significant increase in paw angle variability for both WT and M83 mice after delivery of α-synuclein exosomes. Data represents the mean ± s.e.m. by t-test **P < 0.01, ***P < 0.001 and ****P <0.0001.

An important tenet of the transmission hypothesis in PD is that α-synuclein in recipient cells can be propagated to misfold, promoting disease progression. To test this, we performed intranasal delivery of α-synuclein exosomes to Snca^-/-^mice (α-synuclein knockout mice). The Snca-/- mice given α-synuclein exosomes did not exhibit any abnormal motor impairments at 6 months of age (Fig. 3d, f). DigiGait analysis did not show any functional gait differences between Snca^-/-^mice given control or α-synuclein exosomes (Extended data Fig. 4). These results indicate that the exosomal transmission of Parkinson’s-like pathology requires recipient cells to contain endogenous α-synuclein, supporting the prion-like templating mechanism for PD onset.

To investigate if nasal delivery of α-synuclein exosomes resulted in Parkinson’s-like brain pathology we performed immunohistochemical analysis for brain amyloids at 6 months of age in WT, M83 and Snca^-/-^mice. Phosphorylation of α-synuclein at position 129 promotes fibril formation and has been identified in Lewy bodies of PD brains^29^. We therefore stained brains for both pSer129 α-synuclein and ubiquitin and identified aggregates in multiple regions of the mouse brain after delivery of α-synuclein exosomes. Alpha-synuclein aggregates were found predominantly in the motor cortex, caudoputamen, hippocampus and substantia nigra of both WT and M83 mouse brains (Fig. 4a, b). We also observed positive staining in the brains of WT and M83 mice given α-synuclein exosomes using a conformation-specific antibody that binds with high affinity to filamentous and oligomeric α-synuclein (Fig. 4c). No α-synuclein aggregates were detected in the brains of mice given control exosomes or in Snca^-/-^mice given α-synuclein exosomes. Staining for cell death using an activated caspase-3 antibody in both WT and M83 brains indicated cell death in similar regions that contained α-synuclein aggregates (Fig. 4d). No caspase-3 staining was observed in Snca^-/-^mice treated with α-synuclein exosomes (Fig. 4d). Moreover, after delivery of α-synuclein exosomes to WT mice a reduction of tyrosine-hydroxylase (TH) positive neurons in the substantia nigra was observed, but not in WT mice given control exosomes (Fig. 4e-g). Delivery of α-synuclein exosomes therefore can promote the accumulation of misfolded α-synuclein in the brain, resulting in cell death and pathology similar to that observed in the PD.

**Fig. 4.**
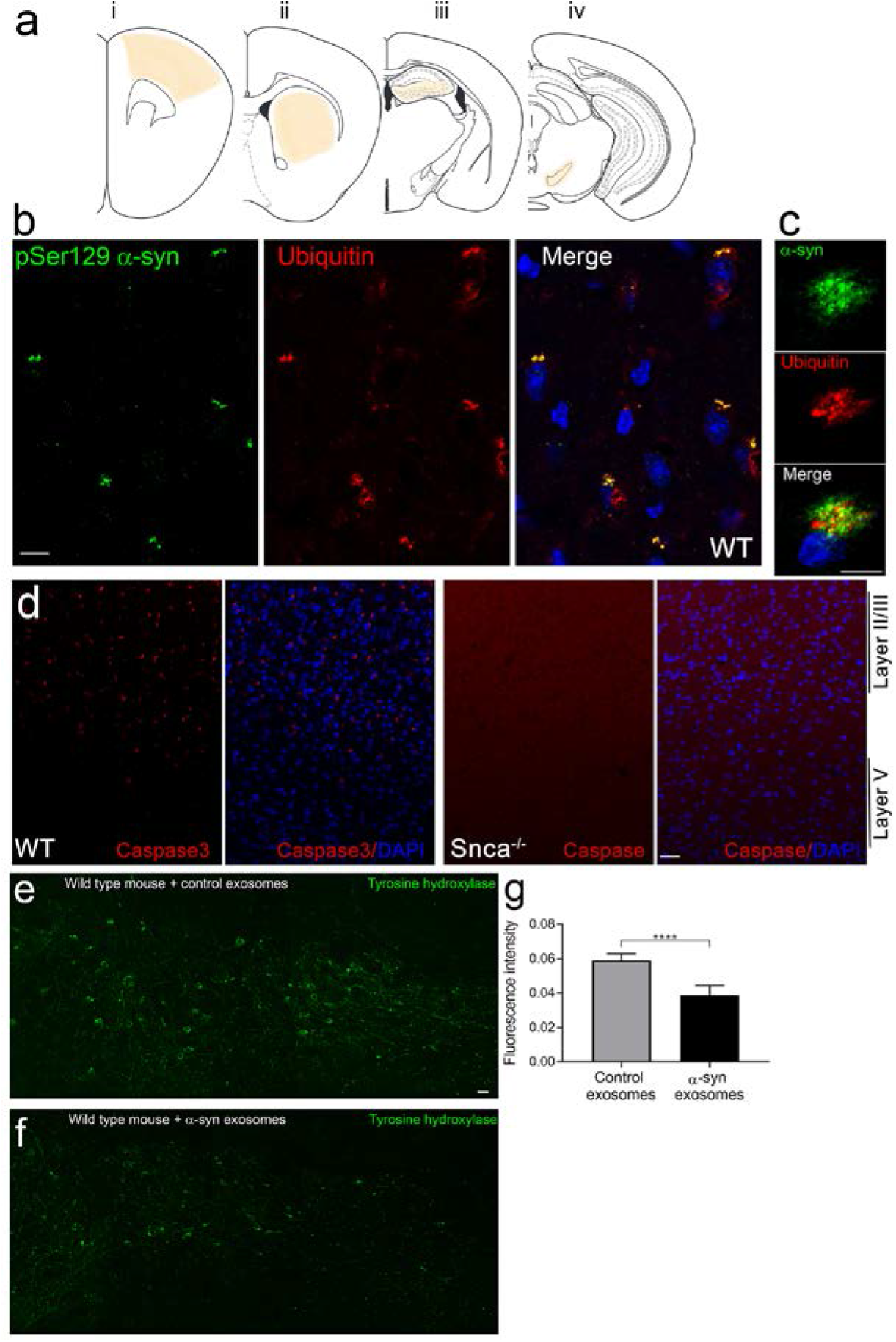
Intranasal delivery of α-synuclein containing exosomes promotes protein aggregates and cell death in the brain. **a**, WT mouse brain after 4 months of weekly intranasal delivery of α-synuclein exosomes contained multiple α-synuclein containing aggregates. The major brain regions that contained pSer129 α-synuclein positive aggregates are shown in peach colour in the schematic. **b**, The motor cortex of a WT mouse after delivery of α-synuclein exosomes showing positive staining for both pSer129 α-synuclein and ubiquitin, indicating brain aggregates. **c**, WT mouse brain after delivery of α-synuclein exosomes stained for a conformation specific α-synuclein antibody (MJFR-14-6-4-2) and ubiquitin showing a magnified view of brain aggregates. **d**, Delivery of α-synuclein exosomes to WT mice resulted in positive staining for Caspase 3 indicating cell death in the motor cortex, delivery of α-synuclein exosomes to Snca^-/-^mice did not result in positive Caspase 3 staining. **e, f**, Immunohistochemical staining for tyrosine hydroxylase positive neurons in the substantia nigra shows a loss of signal in both the cell body and processes in wild type mice given α-synuclein exosomes compared to delivery of control exosomes. g, Quantification of the tyrosine hydroxylase positive signal in the substantia nigra of wild type animals after delivery of exosomes (n =3 mice per group, data represents the mean ± s.e.m. Student T-tests ****P < 0.001). Scale bar; b, c = 10µm, d = 50µm, e, f = 20µm

## Discussion

The majority of PD is idiopathic, and although genetics has been powerful in identifying important pathways linked to the disease, we still have very limited understanding of what causes PD. Here, we provide evidence that α-synuclein carried within exosomes can transmit to recipient brain cells and generate Parkinson’s-like pathology *in vivo*, providing a plausible pathway for the initiation and development of PD. This mechanism is abetted by the Ndfip1/Nedd4-1 pathway, presenting new targets for therapeutic treatment at the earliest time point in the disease progression. Importantly, α-synuclein, Ndfip1 and Nedd4-1 are expressed widely in the body, thus exosomal loading of α-synuclein may occur in multiple regions, including the gut, allowing for a prion-like spread in the body. Our discovery has implications for current immunotherapy approaches to PD treatment; if pathogenic α-synuclein is encased within the exosome microenvironment then antibody based treatments may have limited effectiveness in treating the disease. We suggest that the role of exosomes in spreading α-synuclein is most likely a stochastic process, reliant on cell stress events, the time period of exposure to risk factors, and vesicle clearance mechanisms in the body. As exosomes can be derived and taken up by various cells types, we propose that synucleinopathies in general could be caused by distinct exosome signalling pathways that can traffic α-synuclein between different cell types in the body.

## Supporting information

Supplemental Figues 1-4

## Acknowledgements

We thank Dr Chris Bye for providing the M83 mouse strain; Professor Sharad Kumar for supplying the Nedd4-1 construct. Funding for this research was provided by the National Health and Medical Research Council (NHMRC), APP1164812.

## Author contributions

J.H. conceived the project; J.H., L.H.L, M.B. and J.Y. performed and analysed experiments. J.H., S.G., A.H. and S.S.T provided feedback during the project and wrote the paper and prepared the figures. All authors commented and approved the manuscript.

## Competing interests

The authors declare no competing financial interests.

## Methods

### In vitro tissue culture

HEK-293T and LN18 cells were cultured in DMEM (Invitrogen) containing 10% FCS, 4 mM L-glutamine and 50 μg/ml PenStrep. HEK-293T cells were transiently transfected with appropriate constructs using Effectene Transfection Reagent kit according to the manufacturer’s instructions (Qiagen). Constructs used included; Ndfip1-Flag in pcDNA3, Nedd4-1 in pcDNA3, EGFP-α-synuclein (Addgene construct #40822), EGFP-α-synuclein A53T (Addgene construct #40823), α-synuclein-HA in pHM6.

LN18 cells were treated with PD associated toxins including iron, lysosome inhibiter and pesticide to promote the upregulation of Ndfip1 and the loading of endogenous α-synuclein into exosomes. Cells were treated with either FeCl_2_, ^25^ chloroquine or Rotenone for 20 hours before harvesting the exosomes and cell lysates.

### Bimolecular complementation assays

The coding region of α-synuclein was amplified by PCR and cloned into pBiFC-VC155 (addgene Plasmid 22011) to generate α-synuclein-VC. Constructs for Ndfip1-VN and VN-Ub have previously be reported^16^. BiFC experiments with the expression of Rab constructs cells were fixed 15 h post-transfection as overexpression of Rab GTPases can result in the abnormal accumulation of vesicles in the cell. After 15 h cells were first washed once by 0.1M PB and then fixed in methanol for 10min at -20°C. After fixation, coverslips were incubated for 5 min with 0.1M PB containing DAPI before a final wash in 0.1M PB. Coverslips were then mounted on slides using Mowial. Imaging was performed on a confocal microscope (FV1000; Olympus).

For colocalisation analysis, the Manders overlap coefficient was used to quantify colocalisation. The Manders coefficient describes the fraction of red (M1) or green (M2) channel pixels that colocalise with pixels from the other channel based on intensity. For the colocalisation analyses, the M1 coefficient was used to normalise for the differential expression levels between the Rab-GTPases. Prior to analysis, deconvolution was performed on each z-stack using the Huygens Remote Manager to enhance the resolution and contrast for greater accuracy during analysis. A consistent region representing a cell from a deconvolved confocal z-stack was applied to all z-stacks prior to analysis to ensure consistency in area of analysis. A threshold level, under which pixel values were considered background was manually set and the same threshold applied to all stacks. To calculate the Manders overlap coefficient, the JACoP plugin for ImageJ was used^30^.

### Protein lysate preparation and immunoprecipitation assays

Mouse brain cortices, or HEK-293T and LN18 cells, were lysed in ice cold RIPA buffer (50 mM Tris, pH 7.2, 0.15 M NaCl, 2 mM EDTA, 1% NP40, 0.1% SDS) with complete Mini Protease inhibitor cocktail (Roche Diagnostics) for 20 min at 4°C. Brain homogenates and cell lysates were cleared of insoluble debris by centrifugation at 13,000 rpm for 15 min at 4°C. Protein concentration of lysates was measured using the BIO-RAD DC protein assay according to the manufacturer’s instructions (Biorad). For precipitation experiments, Protein G beads (Pierce) were used for the precipitation of antibodies. Other beads used include Flag beads for Flag tagged Ndfip1. For each experiment, beads were washed 4 times with RIPA buffer before elution using the manufacturer’s instructions.

### Western blotting

Lysates or immunoprecipitates were resolved on 12% SDS-PAGE gels followed by transfer onto Hybond C nitrocellulose membrane (Amersham). After transfer membranes were fixed with 0.4% PFA in PBS for 30 min at RT. Membranes were blocked for 1 h at RT in 5% non-fat milk in TBS, 0.05% Tween-20 (TBST). Blots were incubated overnight with primary antibodies at 4°C followed by appropriate HRP-conjugated secondaries for 1 h at RT. Proteins were detected using Amersham enhanced chemical luminescence reagent as per the manufacturer’s instructions (GE Healthcare) and visualized by exposure to x-ray film.

### Denaturing ubiquitin assay

Ubiquitination assays were performed under denaturing conditions to limit non-specific protein interactions. HEK-293T cells were transfected with the appropriate constructs of His-ubiquitin, α-synuclein, Ndfip1-Flag and Nedd4-1. Twenty-four hours after transfection, cell lysates were prepared and immunoprecipitated with HisLink under denaturing condition using 6M Guanidine hydrochloride and NEM for inhibiting the deubiquitinase enzymes. Beads were washed 3 times before ubiquitinated proteins were eluted using 300mM Imadazole. Eluted fractions were suspended in Laemmli buffer for SDS-PAGE and analysed by Western blotting.

### Isolation of Exosomes

Exosomes were purified either from LN18 cells after treatment with PD associated toxins, or from 293T cells that had been transfected. 293T cells were cultured for up to 72 h post transfection before supernatant was collected. LN18 cells containing endogenous α-synuclein were cultured for 24h after treatment with PD associated toxins before supernatant collection. The supernatant was cleared from dead cells and debris by centrifugation for 10 min at 200g followed by centrifugation for 20 min at 20,000g. Small EVs (containing exosomes) were isolated from the supernatant by centrifugation for 70 min at 100,000g and washed with ice-cold phosphate-buffered saline (PBS) before being centrifuged again for 70 min at 100,000g. Exosomes used for Western blotting were resuspended in 30µl of RIPA buffer and boiled after the addition of 10µl Laemmli buffer before application. Exosomes used for uptake or intranasal delivery were resuspended in HEPES buffer at a protein concentration of ∼2µg/µl. Real-time high-resolution particle detection, counting, and sizing were performed on the NanoSight NS300 following manufacturer protocols (Malvern Instruments, Malvern, UK). Particle concentration (particles/mL) and size was calculated by the NanoSight system.

### Intranasal delivery of EVs to mice

The use of animals in this study conformed to the Australian National Health and Medical Research Council’s published Code of Practice for the Use of Animals in Research and was approved by the Florey Neuroscience Institute animal ethics committee, and by the Swinburne ethics committee. Mice were kept in a 12 h:12 h light:dark cycle at 21-23°C and 30% humidity, with standard mouse chow and water ad libitum. Intranasal delivery of exosomes was performed weekly from 2 months of age. Mice strains used in this study were the A53T mouse^28^ (M83 line) on a C57Bl/C3H background, and SNCA knockout mice on a C57/Blk background. Mice were randomly assigned to each group, and both male and female mice were used for experimentation. Exosomes from LN18 cells purified from three different conditions; (i) control exosomes from unstressed LN18 cells, termed Control exosomes, (ii) Fe spiked exosomes from unstressed LN18 cells that after purification had 400µM FeCl_2_ added, termed Fe spiked exosomes, and (iii) LN18 cells stressed for 20 hours with FeCl_2_ (400µM) before collection of exosomes, termed α-syn exosomes. All exosomes were purified on the day of use for nasal delivery. Purified exosomes were intranasally administered to anesthetised mice. Mice were mildly anesthetised in an induction chamber using 2% isoflurane to sedate the animal. Once sedated, intranasal delivery of exosomes was performed whilst holding the animal in a supine position on a tilted plan with the nose pointing approximately horizontal. Exosomes were delivered from a small tip pipette in a single drop (∼2.5µL) intranasally to one side of the nasal cavity, before waiting 20 seconds and then delivered to the other nasal cavity. After delivery the mouse was returned to the handling box to remain sedated. This procedure was repeated for a total delivery volume of 10µL of exosomes (2µg/µl total protein concentration, approximately 5 × 10^9>^ exosomes delivered). Total time for the intranasal delivery for exosomes was approximately 2-3 minutes per mouse.

### Behavioural analysis

Before each test, mice were habituated in the testing room for 30 min. Behavioural testing of motor function was conducted once per month after nasal delivery was initiated at 2 months of age. Investigators were blinded to the genotype and treatment group for each of the behavioural tests conducted. Tail suspension testing was conducted every two weeks during intranasal delivery of exosomes. Motor function testing included beam walk, pole test and DigiGait analysis of treadmill running and paw angle analysis. All motor function testing was conducted in the late afternoon, close to the beginning of the active period. For all behavioural tests animals were trained in each paradigm at 2 months of age. All animals were able to complete each test without failure at this time.

For beam walk analysis mice were assessed for the number of foot faults on the beam whilst walking over a 60cm distance on a 1.5cm thick rod mounted 15cm above a padded surface. The rod was positioned so the end of the beam led directly into the animal’s home cage. Mice were tested 3 times and the score averaged at each monthly testing session. A score of four indicates no foot faults on the test, a score of three represents <3 foot faults, a score of 2 indicates >3 foot faults and a score of 1 represents freezing on the beam unable to walk.

Balance and motor coordination were assessed by vertical pole test. To perform this task, mice were placed on the top of a pole (60 cm high, 1.5 cm thick), which is fixed vertically to a stable base with small padding on the surface. Each mouse was placed mid-pole facing up, and all four limbs grasp the pole to begin the test. The animals were previously trained to turn on the pole and orient downward to climb to the base. The test was scored on the ability of the mouse to turn on the pole and climb down, representing a fully completed turn on the pole. A score of four indicated a complete turn on the pole within 3 seconds after the turn was initiated, a score of three represented a mouse that found it difficult to turn its hind limb on the pole, taking longer than three seconds to complete the turn once it was started. A score of two represented a mouse that was unable to complete a turn on the pole test. A score of one was a mouse that could not complete the turn and did not have grip strength to hold the pole, resulting in a fall.
Scoring presented is the mean of three trials of the test, with a 3-5 minute break between each test being conducted.

For treadmill running, mice were scored on the ability to run at 30cm/sec for 10 seconds on a 25cm long flat surfaced treadmill. A score of three indicates a mouse that could run at 30cm/sec without falling towards the back of the treadmill, a score of two indicates a mouse that could not run for 10 seconds and had fallen towards the back of the treadmill and a score of one indicated a mouse that could not run at 30cm/sec at all.

For gait analysis of mice, a DigiGait apparatus (Mouse Specifics, Framingham, MA) was used for ventral plane videography of mouse gait kinematics on a moving transparent treadmill belt. Each mouse was trained at three different speeds (15cm/sec, 25cm/sec and 30cm/sec) at 2 months of age. Mice were tested at these speeds each month thereafter. Analysis of video recordings was performed using DigiGait analysis program.

### Immunohistochemistry

For α-synuclein, ubiquitin and tyrosine hydroxylase immunostaining, mice were killed under deep anesthesia by transcardial perfusion of PBS (pH 7.4) followed by 4% paraformaldehyde in 0.1 M phosphate buffer (PB). Brains were post-fixed for 1 h in 4% PFA in 0.1 M PB and then cryoprotected for 24 h in 20% sucrose in 0.1 M PB at 4°C. Brains were embedded in Tissue-Tek optimum cutting temperature compound and stored at -80 °C before sectioning on a cryostat. Perfusion fixed sections (12 μm) were permeabilized with 0.3% Triton X-100 in 0.1 M PB and blocked with 10% FBS in 0.1% Triton X-100 with 0.1 M PB. Sections were incubated with primary antibodies overnight followed by appropriate secondary antibodies for 1 h at RT. Cell death was detected using Caspase3 staining. Sections were counterstained with DAPI (1:10,000, Dako) before mounting under glass cover slips with anti-fade mounting reagent. Fluorescent images of staining were obtained at RT with a 40X or 63X objective on a laser scanning confocal microscope (Olympus FluoView FV1000 or FV3000) using FV100-ASW software (Olympus).

Tyrosine hydroxylase levels in the substantia nigra (SN) of mice was imaged and then analysed using ImageJ. The total fluorescence intensity was calculated from images of the SN of each mouse brain. Three brain sections per mouse were averaged and three separate mice were used per group.

### Antibodies used in this study

**Antibodies used in this study:**
| Antigen | Company | Product number |
| --- | --- | --- |
| α-synuclein | Cell Signaling | 2628 |
| α-synuclein (phos S129) , EP1536Y | Abcam | ab51253 |
| α-synuclein filament antibody [MJFR-14-6-4-2] | Abcam | ab209538 |
| Ubiquitin, clone Fk2 | MBL Life Science | D058-3 |
| β-actin, clone AC-40 | Merk | A4700 |
| anti-Flag M2 | Merk | F3165 |
| Tyrosine Hydroxylase, clone LNC1 | Merk | MAB318 |
| Tsg-101 | Merk | HPA006161 |
| Ndfip1 | Howitt *et al.*<sup>31</sup> |  |
| GM130, Clone 35/GM130 | BD Biosciences | 610822 |

### Statistics and reproducibility

All experiments were performed with at least three technical replicates (see figures for exact number) with similar results. Statistics were performed using a Prism (GraphPad) software. Data were mean ± s.d. unless otherwise noted. A 95% confidence interval was used to define the level of significance. *P < 0.05, **P < 0.01, ***P < 0.001, ****P < 0.0001; NS, not significant. All other pertinent information, including sample size and statistical test used can be found in the figure legends or labelled within the figure.

## Notes

### Competing Interest Statement

The authors have declared no competing interest.

