## Supplemental Figues 1-4 for "Exosomal transmission of α-synuclein initiates Parkinson’s disease-like pathology"

### Extended Data Figures

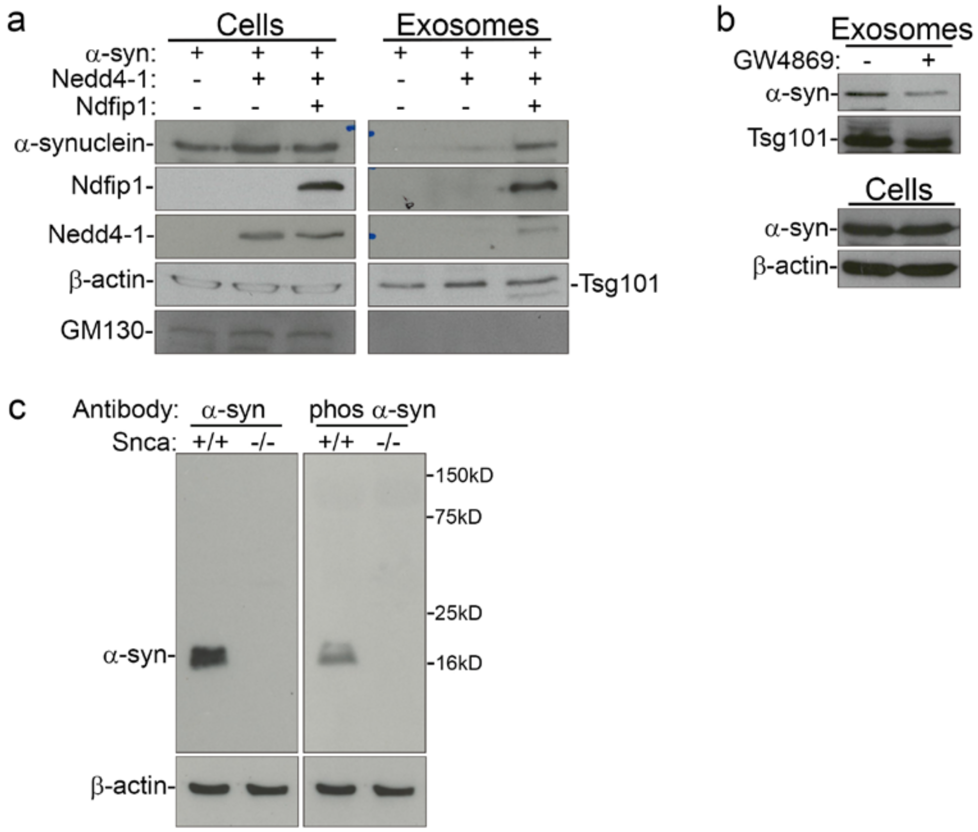


**Extended Data Fig. 1 │ Packaging of α-synuclein into exosomes requires Ndfip1 and can be inhibited by GW4869. a,** Co-expression of α-synuclein with Nedd4-1 does not result in α-synuclein packaging into exosomes. Expression of α-synuclein, Nedd4-1 and Ndfip1 results in the packaging of α-synuclein into exosomes. β-actin is a cell lysate load control, Tsg101 is an exosome load control, GM130 is cellular marker not found in exosomes. **b,** The neutral sphingomyelinase inhibitor GW4869 can prevent the exosomal release of α-synuclein. **c,** Alpha-synuclein antibodies used in this manuscript show detection of a band at ~16.5 kD in a wild type mouse brain cortical lysate that is not present in the Snca$^{-/-}$ mouse brain cortical lysate.

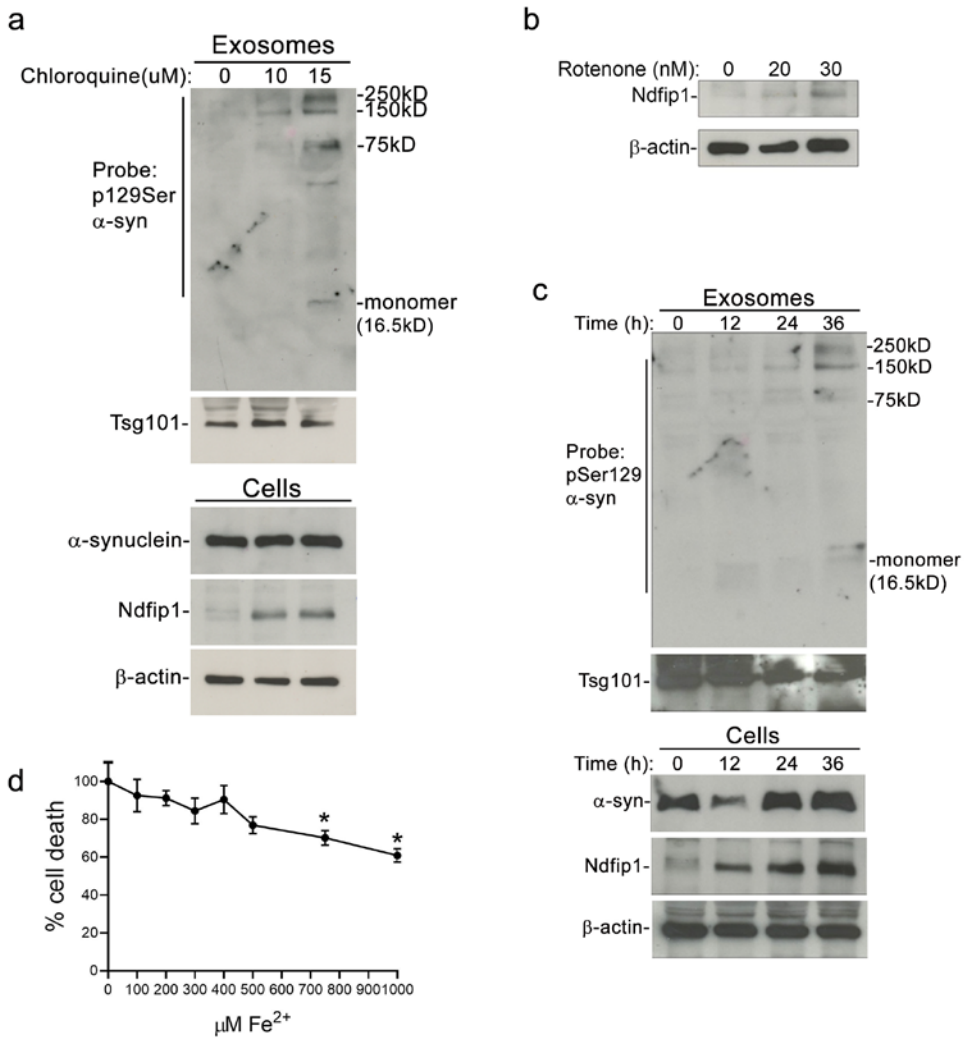


**Extended Data Fig. 2 │ Ndfip1 is upregulated by risk factors for PD, resulting in the loading of α-synuclein into exosomes. a,** Western blot of both cell lysate and exosome preparations shows that increasing concentrations of the lysosome inhibitor chloroquine results in the upregulation of cellular Ndfip1 and the concomitant packaging of α-synuclein into exosomes. **b,** Ndfip1 is upregulated in a dose dependent manner by the pesticide rotenone. **c,** Western blot showing an increase in the aggregation of α-synuclein in exosomes over time. Cells were stressed with 10μM chloroquine for different time periods. **d,** Cell death assay for increasing concentrations of $FeCl_2$ in LN18 cells. Below 500μM there was no significant increase in cell death after 24 hours. Data are the mean ± s.e.m. * $P < 0.05$ from three independent test. One-way ANOVA with Dunnett's multiple comparisons test.

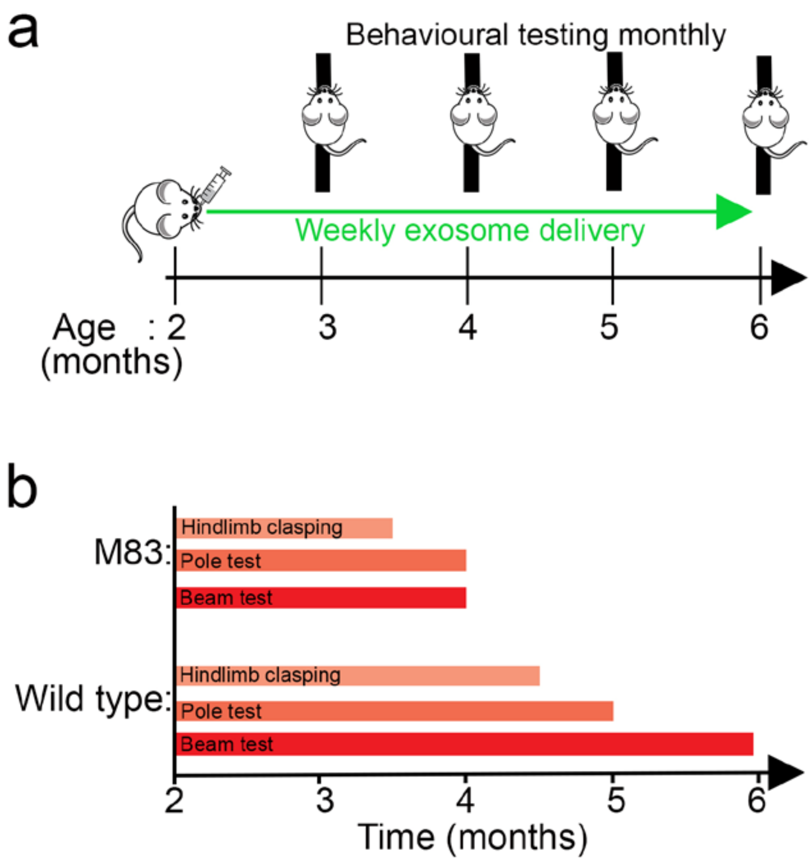


**Extended Data Fig 3 │ Schematics for the delivery of exosomes to mice and the timeline for the onset of motor function deficits. a,** Schematic for the nasal delivery of either control or α-synuclein containing exosomes weekly over a 4 month period, behavioural testing was conducted each month. **b,** Mice were monitored weekly (tail suspension) or monthly (pole and beam test) for the onset of any motor function deficits. Time point for the onset of motor deficits is shown in the figure, deficits were observed earlier in the M83 transgenic mouse compared to wild type mice given α-synuclein exosomes.

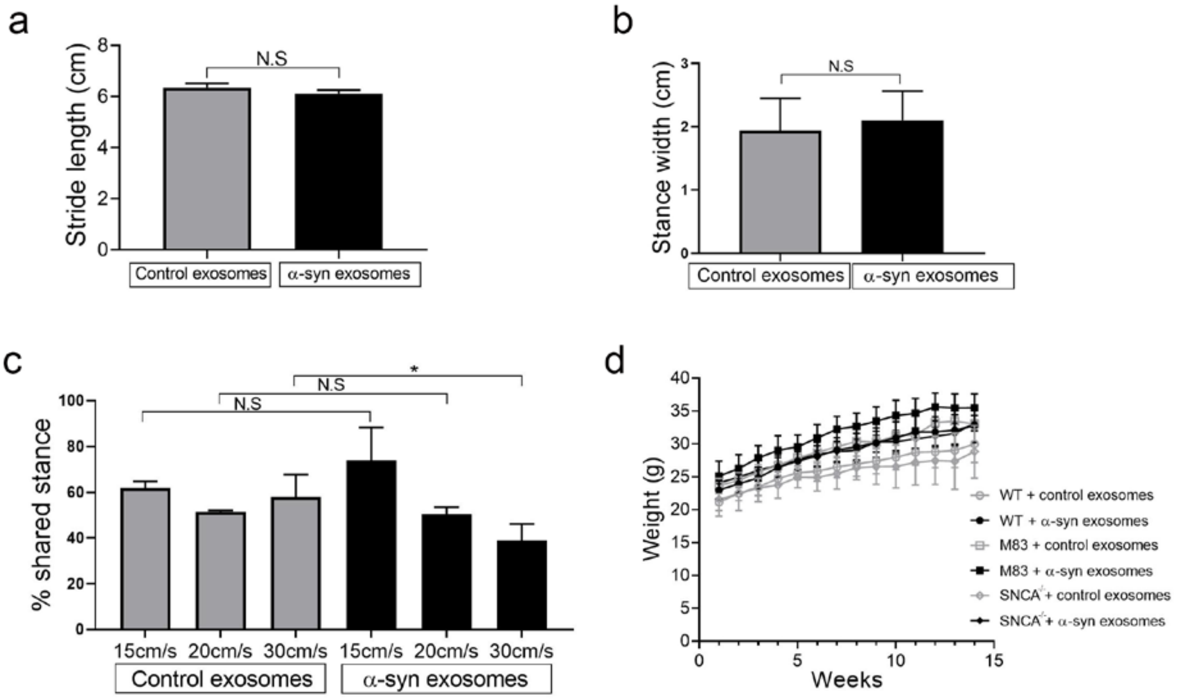


**Extended Data Fig. 4 │ Behavioural analysis of mice after nasal delivery of exosomes. a-b,** Wild type mice give control or α-synuclein exosomes did not show any difference in stride length or stance width metrics using DigiGait analysis. **c,** Wild type mice given α-synuclein exosomes showed a significant difference in shared stance (the percentage of hind limb paws contacting the ground at the same time) at higher running speeds compared to delivery of control exosomes. **d,** No significant difference was observed in weight gain during aging for mice given either control or α-synuclein exosomes across all genotypes tested.
